# Growing *Staphylococcus aureus* in Synthetic Cystic Fibrosis Medium Promotes Colonization in a Murine Pneumonia Model

**DOI:** 10.1101/2025.10.14.682363

**Authors:** Justin M. Luu, Ian N. Moore, Dina A. Moustafa, Joanna B. Goldberg

## Abstract

*Staphylococcus aureus* is a leading cause of bacterial infections worldwide and can lead to diseases such as osteomyelitis, skin, and lung infections in humans. Murine mouse models have been essential to the study of *S. aureus* virulence but are not without their limitations. In murine pneumonia models, colonization of the lungs by *S. aureus* are generally not well maintained. To increase the level and duration of *S. aureus* respiratory colonization, various methods have been employed including embedding the bacteria in agar beads and suppressing the mouse’s immune system. These modifications have improved colonization, but they do not accurately represent clinical infections of diseases such as cystic fibrosis (CF). We hypothesize that culturing *S. aureus* in a different media such as Synthetic CF Medium 2 (SCFM2) would increase colonization compared to growing the bacteria on standard rich media agar plates. We observed that culturing 6 different *S. aureus* strains in SCFM2 led to either a neutral or increased level of lung colonization compared to agar plates. For one strain, WU1, culturing in SCFM2 improved colonization in the oropharynx compared to agar plates and led to a sustained long-term infection in the lungs. Finally, when cultured in SCFM2 compared to agar plates, infection with WU1 led to increased inflammation in both the left and right lung lobe. Overall, we have shown that culturing *S. aureus* in different conditions prior to infection impacts colonization and host response.

**IMPORTANCE:** *Staphylococcus aureus* is a bacterial pathogen that can infect multiple anatomical sites in humans. To study *S. aureus* virulence, murine mouse models have been an essential tool. However, it has been difficult to establish and maintain these infections. To improve *S. aureus* pneumonia murine models, changes to the bacterial dose and mice have been utilized but limits its accuracy to represent clinical infections. In this study, we compared how culturing *S. aureus* in two different conditions prior to infection impacts colonization. We found that the effects of growing *S. aureus* in these different media on acute infections is strain specific. Following up with one of these strains, we showed that Synthetic Cystic Fibrosis Medium 2 (SCFM2) increases *S. aureus* colonization in the oropharynx and leads to a sustain long-term infection in the lungs. Additionally, we showed how culturing *S. aureus* in these conditions impacts the host response.

## INTRODUCTION

*Staphylococcus aureus* is a Gram-positive pathogenic bacterium and a leading cause of bacterial infections worldwide (1). This bacterium can colonize a variety of mammals including humans (2). In humans, *S. aureus* can persist in multiple anatomical sites and can cause a wide variety of infections such as osteomyelitis, endocarditis, skin, and lung infections (3). Additionally, approximately 30% of the human population is asymptomatically colonized with *S. aureus* in their nasal cavity (4).

Murine models, particularly mice, have been an essential tool to study *S. aureus* in the context of infections (5). They offer many advantages such as small size, relative ease of breeding, cost efficiency, a well-characterized immune system, as well as the ability to generate transgenic strains. For these reasons, mice have been used to study infections induced by *S. aureus* such as sepsis, osteomyelitis, endocarditis, and pneumonia (6–9). However, in murine pneumonia infection models, there are difficulties with maintaining bacterial colonization which leads to issues with assessing potential treatments (10). When testing *S. aureus* strains in mice in the context of lung infections, most studies have had to administer supraphysiological (10^7^-10^8^ CFUs) doses intranasally to mice to establish infection (9–13); doses below this threshold generally result in clearance rather than colonization.

To improve the sustainability of *S. aureus* during pneumonia infections in mice, modifications to the preparation of the bacterial dose and the strains of mice used have been explored. Administration of *S. aureus* embedded in agar beads intrathecally to mice have led to prolonged infections that were sustained for at least 21 days (14, 15). Cyclophosphamide, an immunosuppressive chemotherapy that leads to the depletion of neutrophils, has also been used to render mice immunocompromised and more susceptible to *S. aureus* infections (16–18). Additionally, changing the choice of bacteria used in studies have been evaluated. The use of mouse-derived *S. aureus* strains has been shown to more persistently colonize the nasopharynx compared to human-derived *S. aureus* strains (19, 20). These approaches have been greatly beneficial to advancing *S. aureus* infection models; however, these modifications have important limitations which have restricted their adoption. In particular, while embedding *S. aureus* in agar beads improves infection sustainability, it does not accurately represent a clinical infection for diseases such as cystic fibrosis (CF).

CF is a multi-organ genetic disease that impacts more than 120,000 people worldwide (21). This disease is caused by mutations in the CF transmembrane regulator, a sodium and bicarbonate ion channel, which negatively impacts mucus clearing in the lungs and leads to the buildup of sputum (22). Microbial pathogens can utilize sputum as a nutrient source to establish and maintain chronic infections (23). As a result, CF patients are frequently infected with respiratory pathogens throughout their life (24). These chronic bacterial respiratory infections remain the leading cause of morbidity in these patients. *S. aureus* is the most frequently isolated bacteria from respiratory samples of CF patients (8) and persistently colonizes these patients (25, 26). Thus, models to study *S. aureus* in the context of lung infections are paramount to understanding its impact in CF infections.

Here, we studied the growth of *S. aureus* in two different culture conditions prior to infection and monitored its impact on murine colonization and host response. Traditionally, to prepare *S. aureus* for infection, bacteria are cultured in rich laboratory media (such as tryptic soy or lysogeny) on plates or in broth prior to infection. Rich media is relatively inexpensive, widely used and studied, and offers consistency between laboratories. However, this growth condition does not replicate the nutrient composition of a human environment. To address this issue, a defined laboratory medium that mimics the nutrient composition of the sputum of CF patients was developed called Synthetic CF Medium 2 (SCFM2) (27). A previous study by Ibberson and Whiteley revealed that when comparing the gene expression of *S. aureus* cultured in rich media to human CF sputum, the rich media transcriptomes clustered independently of human samples (28). Further they observed that when cultured in SCFM2, the *S. aureus* transcriptomes were more similar to *S. aureus* from CF sputum compared to rich laboratory media (28). Here, we propose that the choice of environmental conditions of *S. aureus* growth prior to infection may improve *S. aureus* colonization during murine pneumonia models and hypothesize that culturing *S. aureus* in SCFM2 prior to infection will increase lung colonization.

To test this, we compared the effects of culturing 6 different *S. aureus* strains using rich media (lysogeny broth; LB) agar plates and in SCFM2 on lung colonization in immunocompetent mice. LB is a commonly used rich laboratory media composed of tryptone, yeast extract, and sodium chloride. As mentioned, SCFM2 contains components meant to mimic sputum from patients with CF; these include mucin, DNA, and amino acids. We chose strains from both human and mouse infections as well as strains from different clonal complexes (CC). *S. aureus* isolates are grouped into different CCs based on their similarity of specific housekeeping genes. In the United States, CC8 and CC5 are the dominate strains isolated from patients and we included strains to represent both (29). We observed that while the effects are strain specific, SCFM2 either had a neutral impact or increased the strains’ ability to colonize the murine lung. In one *S. aureus* strain known to persistently colonize the oropharynx in mice, WU1, we found that culturing in SCFM2 improved colonization levels in the oropharynx. We also discovered that culturing WU1 in SCFM2 leads to sustained colonization in the lungs for at least 4 days. Furthermore, infections with this strain cultured in SCFM2 led to increased inflammation and altered cytokine levels in the left and right lung lobes compared to growing this strain on agar plates. Overall, we have shown the impact of culturing *S. aureus* in SCFM2 compared to agar plates on colonization and host response in a murine pneumonia model.

## RESULTS

### *S. aureus* strains used in this study exhibit both clinical and genomic diversity

The goal of this study is to characterize how altering culture conditions impacts *S. aureus* colonization during murine pneumonia. To achieve this, we sought to test a diverse set of strains under two growth conditions. We used 6 *S. aureus* strains: 5 strains isolated from human infections and 1 strain isolated from a mouse infection (Table 1). JE2 and Newman, both CC8 strains, are frequently used as model strains for methicillin-resistant (MRSA) and methicillin-sensitive (MSSA) strains, respectively (30, 31). JE2 is a derivative of a strain isolated from a skin abscess, while Newman was isolated from a tubercular osteomyelitis infection. N315, an MRSA pharyngeal isolate, was chosen as a representative CC5 strain (32). Sa_CFBR_43 (MRSA) and Sa_CFBR_46 (MSSA) was isolated from respiratory samples of patients with CF (33). Sa_CFBR_43 belongs to CC8 while Sa_CFBR_46 belongs to CC45, a lineage of *S. aureus* that has two classes of *S. aureus* quorum sensor regulators (34). WU1 (MSSA) was isolated from an outbreak of preputial gland infections in a mouse breeding facility and belongs to CC88, a CC that is primarily isolated from rodents in the Western World (20, 35). We determined the diversity of these strains genomically through average nucleotide analysis comparisons (Table S1) (Figure 1). Perhaps not surprisingly JE2, Newman, and Sa_CFBR_43, all being CC8, shared greater than 99.9% similarity. Sa_CFBR_46 was the most genetically distant compared to the other strains followed by WU1.

**Figure 1.**
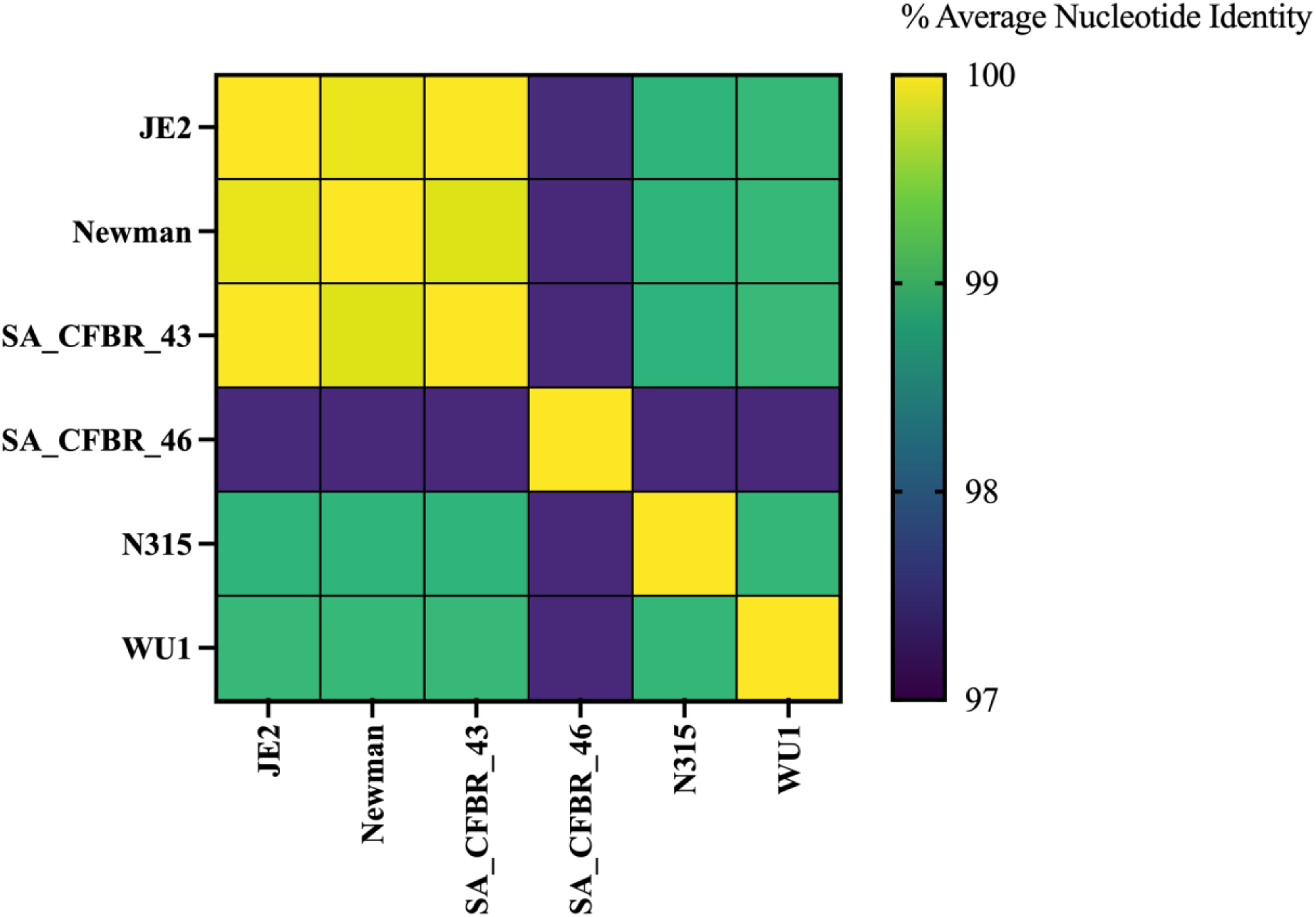
Strains used in this study exhibit genetic diversity. The genomic sequences of each *S. aureus* were compared to each other, and average nucleotide identity was determined utilizing pyani. The numeric results were used to generate the heatmap.

**Table 1.**
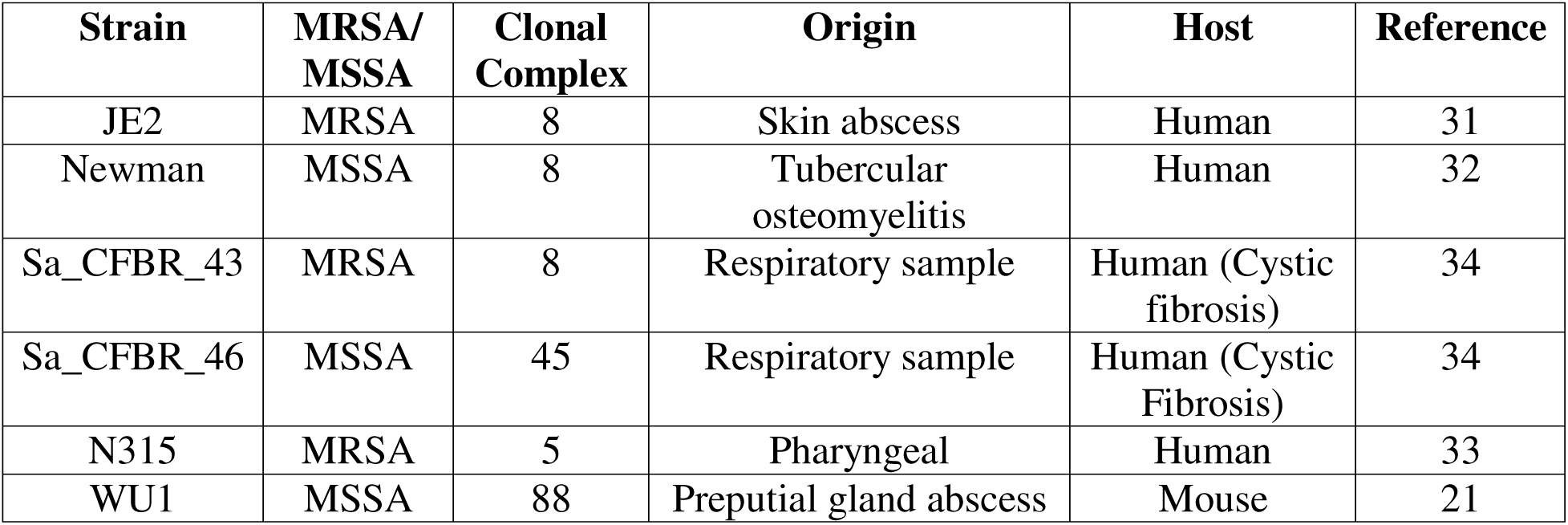
*Staphylococcus aureus* strains utilized in this study.

| Strain | MRSA/<br>MSSA | Clonal<br>Complex | Origin | Host | Reference |
| --- | --- | --- | --- | --- | --- |
| JE2 | MRSA | 8 | Skin abscess | Human | 31 |
| Newman | MSSA | 8 | Tubercular<br>osteomyelitis | Human | 32 |
| Sa_CFBR_43 | MRSA | 8 | Respiratory sample | Human (Cystic<br>fibrosis) | 34 |
| Sa_CFBR_46 | MSSA | 45 | Respiratory sample | Human (Cystic<br>Fibrosis) | 34 |
| N315 | MRSA | 5 | Pharyngeal | Human | 33 |
| WU1 | MSSA | 88 | Preputial gland abscess | Mouse | 21 |

### Culturing *S. aureus* strains in SCFM2 leads to similar growth kinetics compared to culturing in LB

LB and SCFM2 differ in nutrient composition which could have an impact on *S. aureus* growth and physiology (27). To determine whether the difference in nutrient composition impacts growth, all strains were cultured statically in either LB or SCFM2 in 6-well plates and sampled for colony forming units (CFUs) every 2 hours for 8 hours as well as at 24 hours (Figure 2). During the first 8 hours of growth, all strains exhibited similar growth kinetics in both media. At the 24-hour time point, there was equal number of CFUs for all strains cultured in both growth media, except for Newman. Newman had a higher CFUs when cultured in LB (10^8.5^ CFUs) compared to SCFM2 (10^8.2^ CFUs) at 24-hours. These results indicate that the strains investigated here exhibit similar growth kinetics in LB and SCFM2.

**Figure 2.**
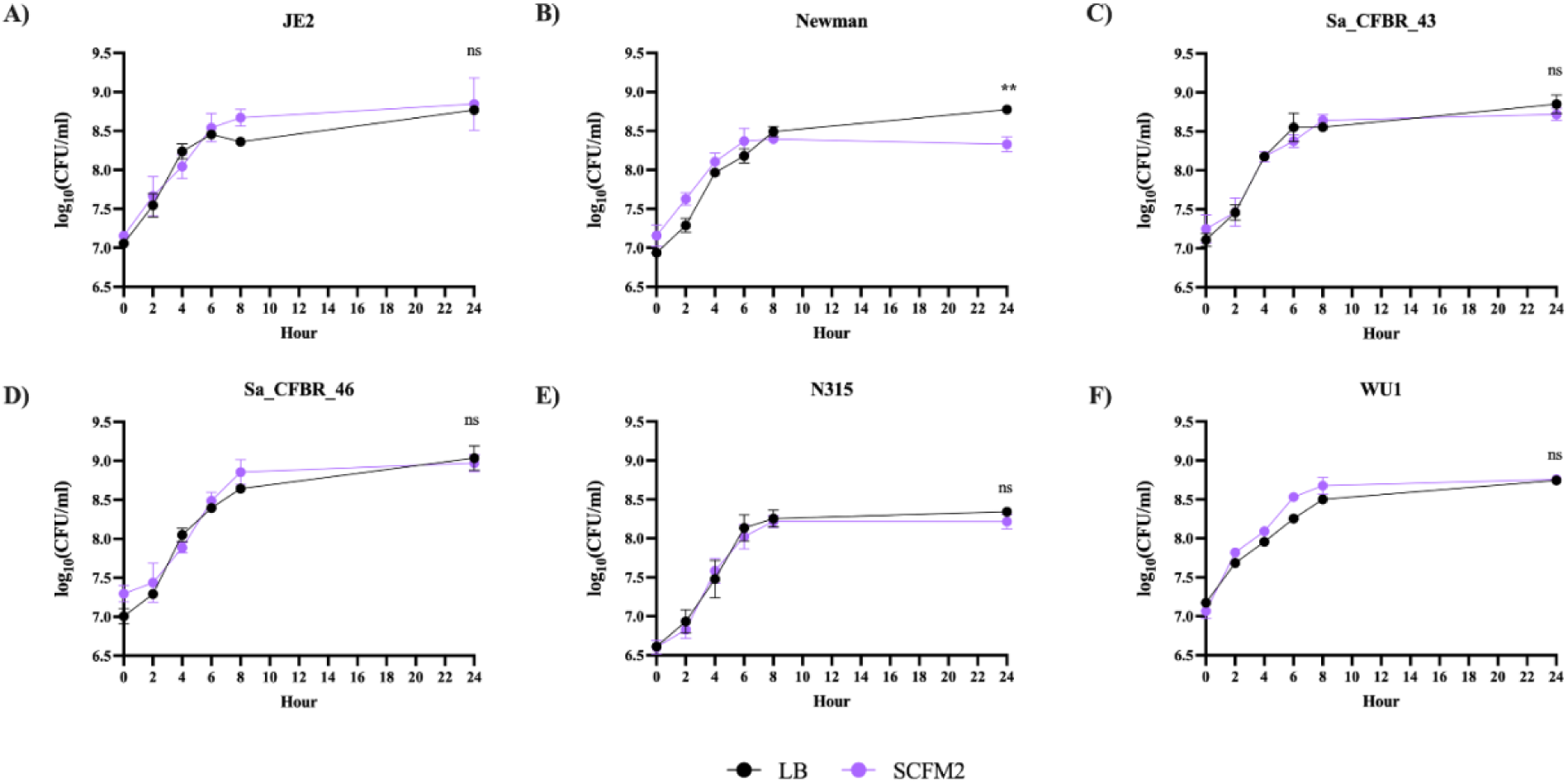
Culturing *S. aureus* in SCFM2 led to similar growth kinetics compared to growth in LB for most strains. Each *S. aureus* strain was grown statically in either LB or SCFM2 in a 6-well plate at a starting OD_600_ of 0.01. Aliquots were removed, serially diluted, and plated on LA plates every 2 hours for 8 hours as well as 24 hours. Each point represents an average of biological triplicates. Statistical analysis was performed on the 24-hour time point through unpaired student *t*-test with Welch correction. **p < 0.01, ns=not significant.

### Culturing *S. aureus* in SCFM2 increases lung colonization in most examined strains

We hypothesized that culturing *S. aureus* in SCFM2 would increase the level of colonization of the mouse lung compared to LB. To test this hypothesis, we infected 8–10-week-old C57BL/6 or Balb/c mice with ∼1 x 10^8^ CFUs of each *S. aureus* strain, prepared through our agar plate and SCFM2 workflow (Figure 3). At 24-hours post-infection, the mice were sacrificed and *S. aureus* colonization in both the lungs (Figure 4) and nasal cavity (Supplemental Figure 1) was determined.

**Figure 3.**
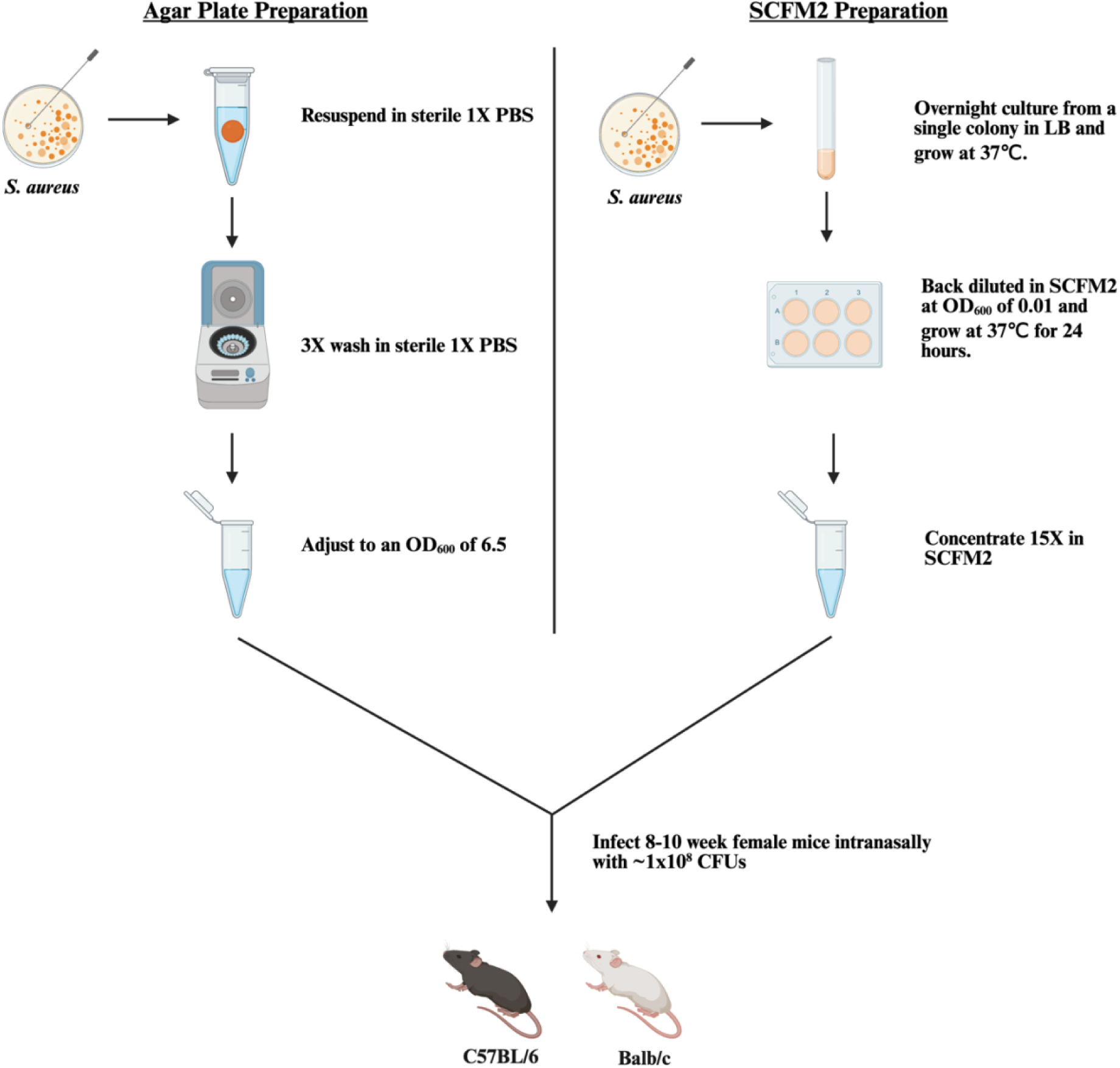
Schematic of the *S. aureus* preparation workflows used in this study. *S. aureus* strains were cultured on either an agar plate (left) or in SCFM2 (right). For the agar plate preparation, *S. aureus* was grown on *Staphylococcus* Isolation Agar (SIA) overnight at 37℃, swabs of colonies were transferred to 1X PBS, washed 3 times in 1X PBS, and adjusted to an OD_600_ of 6.5. For the SCFM2 preparation, a single colony from *S. aureus* grown on SIA was grown in LB overnight with shaking at 37℃. The culture was washed 3 times in 1X PBS, back diluted in SCFM2 at a starting OD_600_ of 0.01 in a 6-well plate, grown statically for 24 hours, and concentrated 15X in fresh SCFM2. 12.5 µl of each condition was subsequently administered intranasally into each nostril (25 µl total) of anesthetized C57BL/6 or Balb/c mice, corresponding to ∼1 x 10^8^ CFUs.

**Figure 4.**
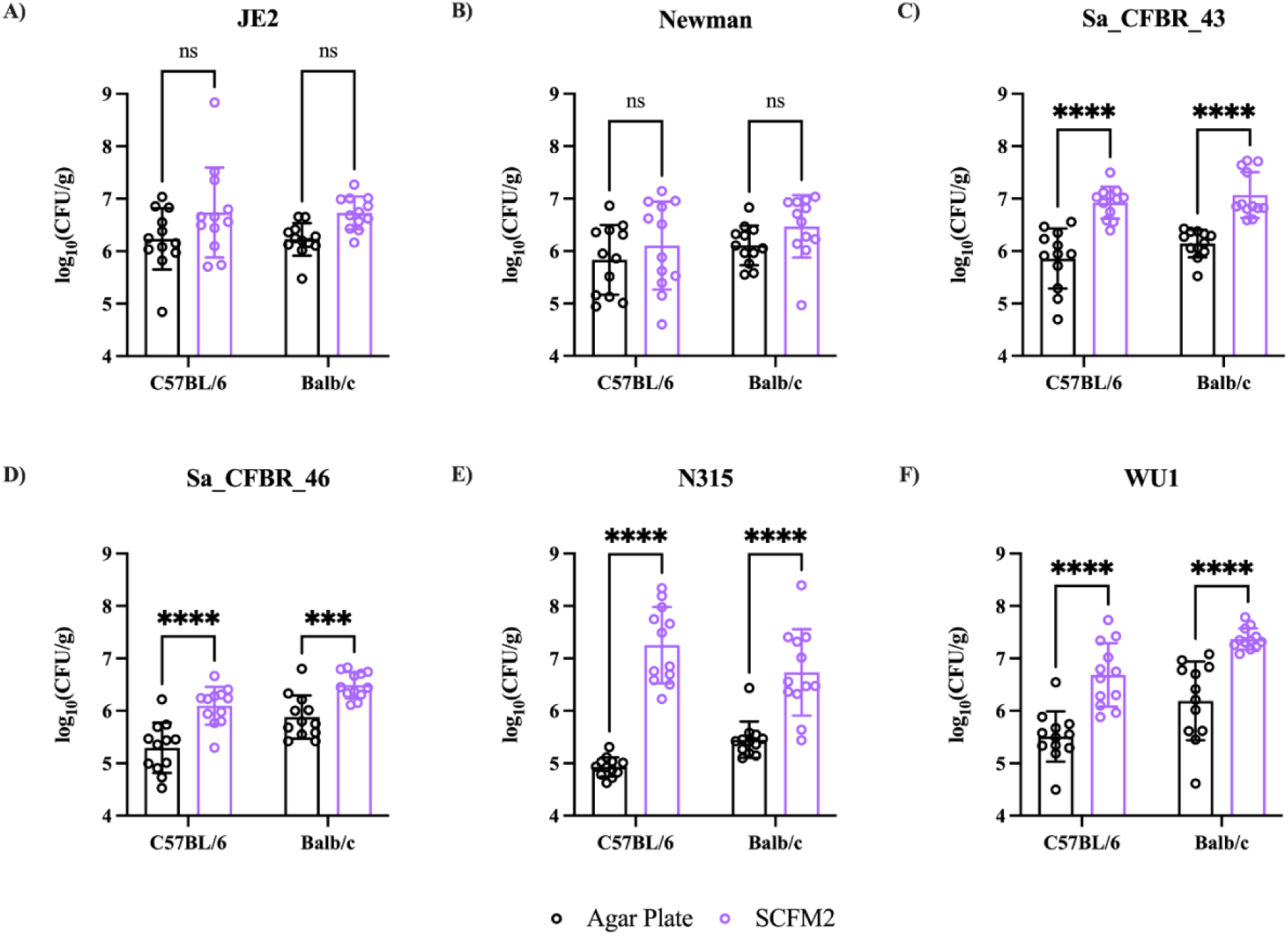
Culturing *S. aureus* in SCFM2 generally increases lung colonization in mice during acute pneumonia independent of mouse strain. *S. aureus* strains were cultured on an agar plate or in SCFM2 as described in Figure 3 and administered to either 8–10-week-old female C57BL/6 and Balb/c mice. Mice were sacrificed at 24 hours post-infection, the lungs were aseptically removed and weighed, homogenized in 1X PBS, and plated on SIA. Each data point represents a mouse. The mean and standard deviation are represented by the error bars. Data were analyzed using two-way ANOVA with Šídák correction. ***p < 0.001, ****p < 0.0001, ns=not significant.

Our results revealed that culturing strains Sa_CFBR_43, Sa_CFBR_46, N315, and WU1 in SCFM2 significantly increased lung colonization in both C57BL/6 and Balb/c mice compared to culturing on agar plates (Figure 4C, 4D, 4E, 4F). In contrast, there was no significant difference in colonization for JE2 and Newman between the culture conditions in either mouse strain (Figure 4A, 4B). When cultured in SCFM2, N315 had the highest average colonization in C57BL/6 mice while WU1 had the highest average colonization in Balb/c mice. The effects of SCFM2 on nasal cavity colonization were more variable (Supplemental Figure 1). While there were no significant differences in JE2 colonization in either mouse strains, Newman had significantly lower colonization in the nasal passage of Balb/c mice when cultured in SCFM2 compared to agar plates while no significant differences in C57BL/6 mice. Both CF isolates, Sa_CFBR_43 and Sa_CFBR_46, had significantly lower colonization in the nasal cavity of either mouse strain. Finally, N315 and WU1 had significantly higher colonization in the nasal cavity independent of mouse strains. Overall, our results indicate that while the effects of growing *S. aureus* in SCFM2 on lung and nasal cavity colonization is strain-specific, it can improve colonization levels in the lungs.

### Culturing WU1 in SCFM2 increases recovery from the throat of mice 2 days post infection and from the lungs at 4 days post infection compared to LB

Since we have observed that the effects of SCFM2 on *S. aureus* colonization were strain specific, we used *S. aureus* strain WU1 as a model system for subsequent studies. WU1 had the highest and most consistent level of colonization in Balb/c mice when cultured in SCFM2, therefore, we chose to concentrate our subsequent analysis on WU1 and Balb/c mice. Previous studies have only characterized WU1’s ability to persistently colonize mice oropharynx during respiratory infections (20, 36, 37) and we aimed to determine if SCFM2 can improve the level and duration of both the throat and lung colonization. We monitored colonization in both the throat and the lungs of Balb/c mice over time after they intranasally infected with ∼1 x 10^8^ CFUs of WU1 grown using either culture condition (Figure 5). In one cohort of mice, throat colonization was determined by swabbing the oropharynx of the same mice on 1, 2, 3-, 7-, 10-, and 14-days post infection (Figure 5A). There was significantly higher oropharynx colonization by WU1 when grown in SCFM2 compared to on agar plates at day 1 and 2 post infection. No significant differences were observed on days 3, 7, 10, and 14 post infection. This suggests that the initial improvement in WU1 oropharynx colonization provided by SCFM2 occurs within 2 days post-infection. Next, we monitored WU1 colonization in lungs over time using separate cohorts of Balb/c mice (Figure 5B). Every day for 4 days, colonization levels in the lungs of the mice were determined. We observed significantly higher and consistent lung colonization up to 4 days post infection when WU1 was grown in SCFM2 compared to agar plates. Overall, our results indicate that, in addition to increasing colonization during early infections, culturing WU1 in SCFM2 increases oropharynx colonization 2 days post infection and leads to sustained lung colonization for 4 days post infection.

**Figure 5.**
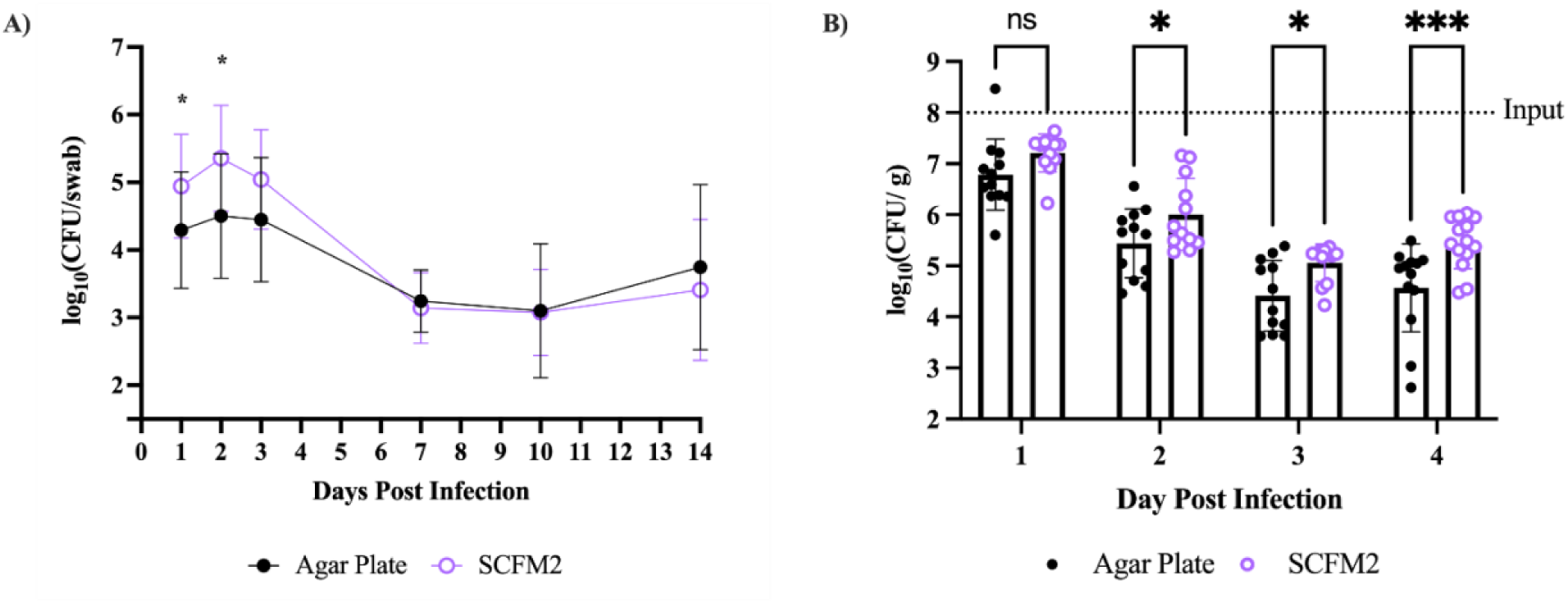
Compared to agar plate preparation, culturing WU1 in SCFM2 improves colonization for the first 2 days post-infection in the throat and for 4 days post-infection in the lungs. *S. aureus* strain WU1 was cultured on an agar plate or in SCFM2 as described in Figure 3 and were administered to 8–10-week-old female Balb/c mice. **A)** The throats of the same mice were swabbed on 1, 2, 3-, 7-, 10-, and 14-days post infection, the swabs were resuspended in 1X PBS, serially diluted, and plated on SIA. Each point represents the average and standard deviation of 12 to 15 mice. **B)** A separate cohort of mice were sacrificed after 1-, 2-, 3-, and 4-days post-infection, the lungs were aseptically removed and weighed, homogenized in 1X PBS, and plated on SIA. Each data point represents a mouse. Statistical analysis for the throat swabs was performed on the 1- and 2-day time point through unpaired student *t*-test with Welsch correction. Multi-day lung colonization was analyzed using two-way ANOVA with Fisher’s LSD post-hoc test. *p < 0.05, ***p < 0.001, ns=not significant.

### Infections with WU1 cultured in SCFM2 leads to increased inflammation and altered cytokine profile compared to agar conditions

We next sought to determine how culturing WU1 in SCFM2 vs. agar plates impacts the host response through histology and cytokine analysis in our pneumonia model. Balb/c mice were intranasally infected with ∼1 x 10^8^ CFUs of WU1 prepared through either culture methods; 1X PBS, and SCFM2 alone served as “vehicle controls”. For these studies, mice were sacrificed 24-hours post infection, and the bacterial load in each lung lobe was determined. We observed a relatively even distribution of CFUs in the left or right lung lobes of mice infected with WU1 cultured in SCFM2 (Supplemental Figure 2B). In WU1 cultured from agar plates, we observed slightly higher CFUs recovered from the left compared to the right lung lobes. Overall, we observed higher levels of colonization in both lung lobes when infected with WU1 cultured in SCFM2 compared to agar plates.

We wanted to determine how WU1 growth conditions prior to infection impacted the response in the lung. To do this, we used two parallel sets of mice for histology and cytokine analysis (Supplemental Figure 2A). In the first set, the left lung lobe was processed for histology while the right lobe was processed for cytokine analysis. In the other set, the opposite lobe was processed for histology and cytokine analysis. Thus, for all these comparisons, the left and right lungs were from different mice.

We assessed the histopathological changes (Figure 6) and cytokine levels (Figure 7) in each lung lobe following infection with WU1 cultured in SCFM2 vs. on agar plates. Overall, we did not observe a major difference between the left and right lobes under any of the conditions tested. First, we monitored the effects of the “vehicle controls” and found that administration of 1X PBS led to scores within the normal histological limits while SCFM2 led to the recruitment of primarily lymphocytes (Figure 6A, 6B, 6E, 6F). These results of the vehicle control were used as the baseline when assessing the histology of the infected lungs. While WU1 cultured from either SCFM2 or agar plates led to inflammation composed of PMN cells, WU1 cultured in SCFM2 led to a higher degree of inflammation as seen by the multifocal and coalescing areas of PMN cells in both lung lobes (Figure 6C, 6D, 6G and 6H). Inflammation of lungs infected with WU1 cultured from agar plates was scored as moderate to severe while lungs infected with WU1 cultured in SCFM2 was scored as severe.

**Figure 6.**
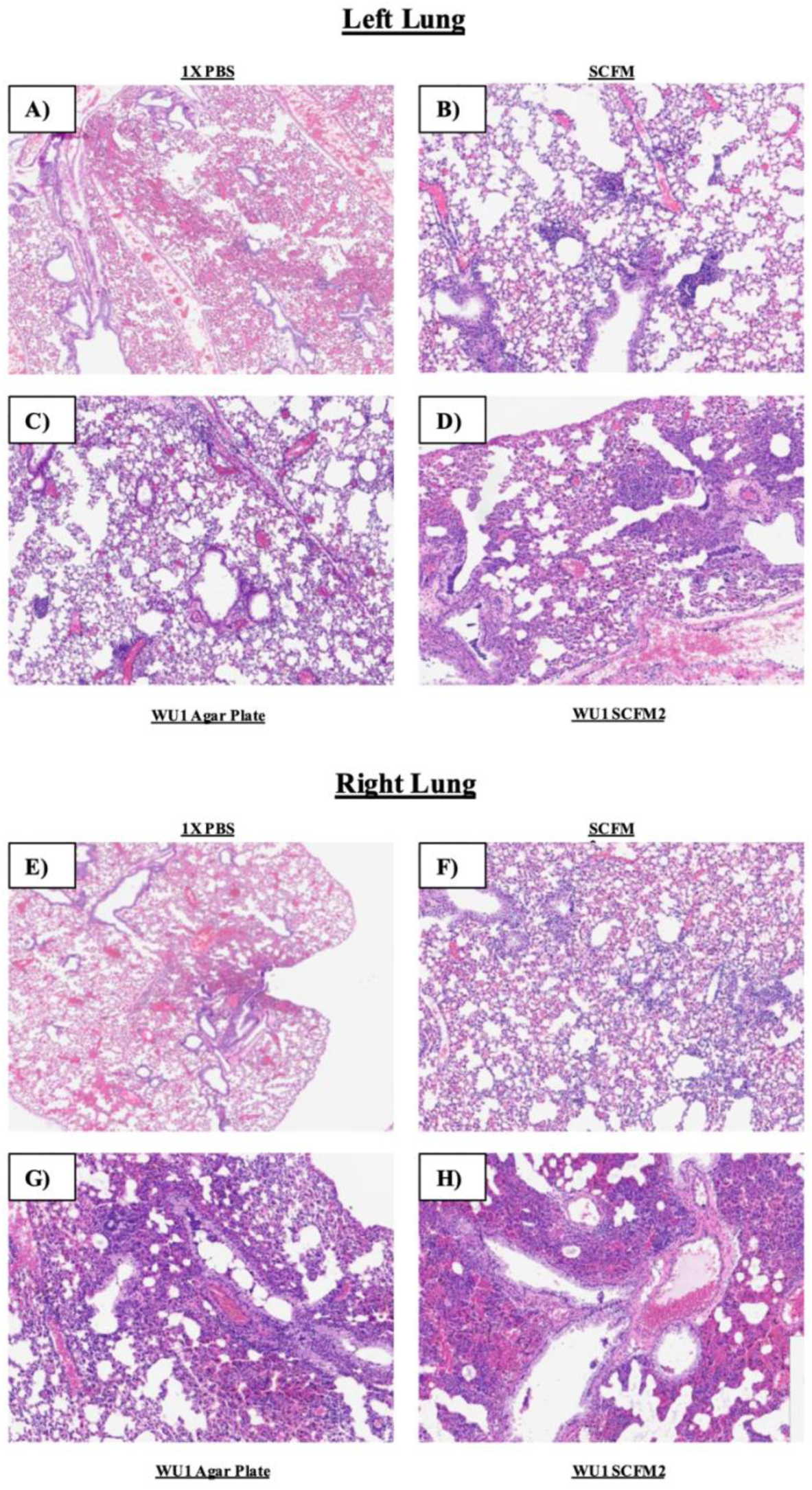
Infections with WU1 cultured in SCFM2 leads to increased inflammation and neutrophil recruitment in both lung lobes compared to agar plate preparation. *S. aureus* strain WU1, cultured either on an agar plate or in SCFM2 as described in the workflow in Figure 3, was administered to 8–10-week-old female Balb/c mice. Mice were euthanized 24 hours post infection. For each culture workflow (Supplemental Figure S2) the left lung **(A-D)** or right lung **(E-H)** lobes were from each mouse (n=6) were inflated and fixed using 4% paraformaldehyde. The lungs were then processed, strained with H&E, and blindly assessed by a veterinary pathologist.

**Figure 7.**
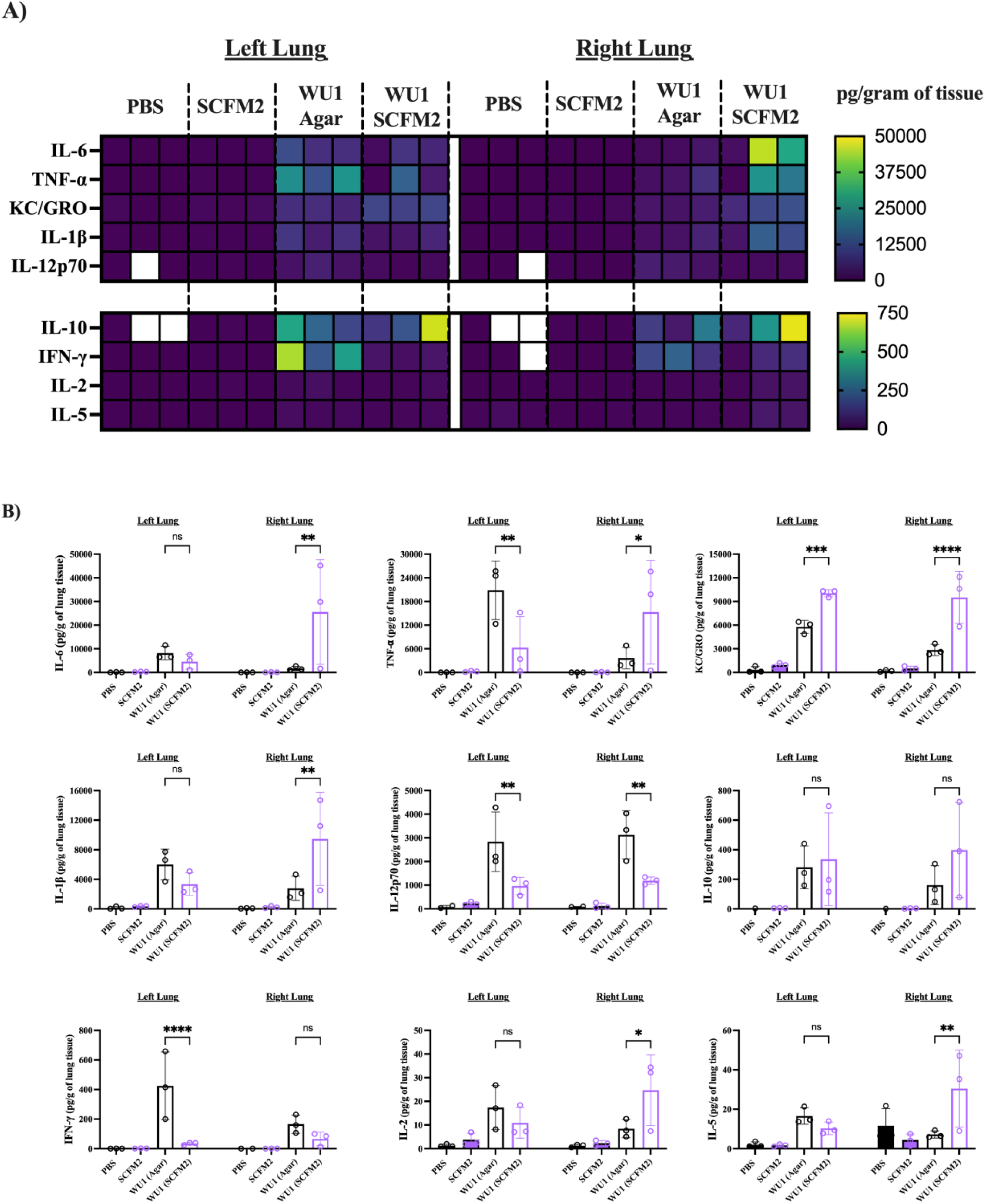
Cytokine profile from lungs infected with WU1 cultured on agar plates and SCFM2. *S. aureus* strain WU1 cultured either on an agar plate or in SCFM2 as described in Figure 3, was administered to 8–10-week-old female Balb/c mice. Mice were euthanized 24 hours post infection. For each culture workflow (Supplemental Figure S2), the left lung or right lung lobes were excised from each mouse (n=6). The lungs were then homogenized in 1X PBS + 1X protease inhibitor, the cell free lysates were isolated, and cytokines were measured with the V-plex proinflammatory cytokine panel 1 (MSD). Quantification of each cytokine was normalized to weight of the lungs and visualized in a heatmap (**A**). Proinflammatory cytokine profiles for lungs infected with WU1 cultured in both conditions were compared (**B).** The mean and standard deviation are represented by the error bars. Statistical analysis was performed using two-way ANOVA with Fisher’s LSD post-hoc test. *p < 0.05, **p < 0.01, ***p < 0.001, ****p < 0.0001, ns=not significant.

Next, we measured the cytokine levels in the other lung lobe. Overall, in both lungs, we generally observed higher cytokine levels in lungs infected with WU1 under either growth conditions compared to the vehicle controls (Figure 7). The only exception was that there was slightly more IL-5 detected in the right lung that was administered the PBS control compared to WU1 cultured on agar plates, however all these levels were very low. In looking specifically at the right lungs, we observed significantly higher levels of the cytokines IL-6, TNF-⍰, KC/GRO, IL-1β, IL-2, and IL-5 in the mice infected with WU1 cultured in SCFM2 compared to agar plates (Figure 7B). On the other hand, IL-12p70, IL-10, or IFN-γ had lower or no significant difference between conditions. For the left lungs, significantly higher levels of KC/GRO were detected in the lungs infected with WU1 cultured in SCFM2 compared to agar plates. TNF-⍰, IL-12p70, and IFN-γ were significantly lower between these conditions while there was no significant differences between the levels of IL-6, IL-1β, IL-10, IL-2, and IL-5. Overall, we observed a cytokine profile that does correlate with the higher levels of inflammation and neutrophil recruitment observed in the histology between the culture conditions. Our findings indicate that culturing WU1 in SCFM2 leads to increased inflammation in both lung lobes that is reflected both in histopathology and cytokine levels.

## DISCUSSION

Laboratory mice have often been used as an *in vivo* model to study *S. aureus* infections such as bacteremia, sepsis, pneumonia, and many others (5). These models have been essential in advancing our understanding of *S. aureus* virulence and host response but have important limitations. When studying *S. aureus* induced pneumonia in murine models, the level of colonization is not well sustained (9, 13, 38, 39). Groups have explored the use of different concentrations of inoculum to induce *S. aureus* pneumonia infection. Parker et al. administered 1 x 10^7^ CFUs of *S. aureus* strain USA300 intranasally to C57BL/6J mice and observed a recovery of 0.5 x 10^7^ CFUs from lung homogenates after 24 hours (38). Torres et al. observed a similar recovery after administering a lethal dose of 3.78 x 10^8^ CFUs of *S. aureus* strain Newman via the same route to C57BL/6J mice after 18 hours (13). Finally, Urso et al. recovered 5 x 10^5^ CFUs of Newman after 24 hours following administration of 1 x 10^8^ CFUs intranasally to C57BL/6 mice (39). Additional modifications include using mouse-adapted strains, altered inoculum preparation methods, and depletion of the mouse’s innate immune response (9, 14, 16–19, 40). While these methods have resulted in infections, the levels of colonization could be improved, especially in immunocompetent mice. Here, we hypothesized that culturing *S. aureus* in a media to mimic the lung environment (SCFM2), would increase colonization.

Our study reveals that improving *S. aureus* colonization during a murine respiratory infection relies on the interactions between several factors: the choice of *S. aureus* strain, the choice of mouse strain, and the growth conditions of the bacteria inoculum. Of the 6 *S. aureus* strains tested, we observed the greatest improved of colonization in N315 for C57BL/6 mice and WU1 for Balb/c mice. WU1’s sustained colonization could be attributed to it being isolated from a mouse and may already be more adapted to evade this specific host response. In mice, culturing JE2 and Newman, two commonly studied *S. aureus* strains, in SCFM2 did not improve colonization in mice while all other strains (Sa_CFBR_43, Sa_CFBR_46, N315, and WU1) showed increased levels of colonization. Except for WU1, all *S. aureus* strains tested were isolated from human infection but only strains isolated from respiratory infections (Sa_CFBR_43, Sa_CFBR_46, and N315) had increased colonization levels in SCFM2.

Interestingly, of the 3 CC8 strains tested, only Sa_CFBR_43 had a significantly higher lung colonization in both mouse strains when cultured in SCFM2 compared to agar plates despite being highly similar genomically to JE2 (99.99%) and Newman (99.85%). In future studies, exploring the genetic differences between JE2 and Sa_CFBR_43 might reveal why SCFM2 increases Sa_CFBR_43 colonization in mice. Similar studies carried out by Sun et al. revealed that the *Staphylococcal* protein A (spA) is required for persistent colonization in the nasal passage of WU1 (20). JE2 and Sa_CFBR_43 share genomically identical sequences for spA but whether differences in expression of this protein contributes to colonization was not investigated in our study. While more strains must be tested, adaptations to the infection site where the strains were isolated from might influence the colonization of *S. aureus*. When developing models for infection and choosing a strain to test, we recommend testing strains isolated from different infection sites as that might impact colonization.

When assessing how different culture conditions impacts colonization of the 6 *S. aureus* strains, we utilized both C57BL/6 and Balb/c mice. These two strains are the most commonly used inbred mouse strains in the field, are the genetic backbone of many transgenic strains, and allow the study of different innate immune responses. C57BL/6 mice exhibit a stronger Th1 immune response while Balb/c exhibit a stronger Th2 immune response (41, 42). Previous studies of WU1 have primarily focused on its ability to persistently colonize the nasopharynx in C57BL/6 mice (20, 36, 37). Our study characterized the ability of WU1 to colonize the lungs of Balb/c mice not only during an acute short-term infection but also during a longer-term infection. Compared to infections in C57BL/6, we found that WU1 colonizes the lungs of Balb/c mice at a higher and more consistent level. We propose that Balb/c mice may be the preferable choice to study WU1 in the context of lung colonization in SCFM2. Additionally, our findings compare the colonization distribution, histology, and cytokine recruitment between the left and right lobes of mice infected with *S. aureus*. The distribution of *S. aureus* in one or another mouse lung lobe following intranasal administration has not been previously studied according to our literature search. We found that WU1 had similar levels of colonization in both the left and right lung lobe as well as similar histopathology scores but a different proinflammatory cytokine profiles. When cultured on agar plates, we observed that WU1 colonization in the left lung was slightly higher than the right lung while the distribution of WU1 cultured in SCFM2 was more even between lobes. Similar pathohistological scoring was between the right and left lungs when infected with WU1 cultured in both conditions. Previous studies had shown that *S. aureus* induced pneumonia in mice leads to inflammation and the recruitment of polymorphonuclear (PMN) cells and our results were consistent with those findings (9, 43, 44). While both culture conditions resulted in inflammation, the degree of inflammation was higher when WU1 was cultured in SCFM2 compared to agar plates. The cytokine profile was generally consistent between the lobes under both culture conditions but there were some variations. Levels of proinflammatory cytokines (IL-1β, IL-6, TNF-⍰, IFN-γ, IL-2, and IL-5) were generally lower in the right lungs compared to the left when infected with WU1 cultured on agar plates but higher when infected with WU1 cultured in SCFM2. In our infection model, there was not a significant difference in WU1 colonization observed but a different host response between the left and right lungs. This suggests that harvesting individual lung lobes may be useful for measuring colonization to decrease the number of mice used but the whole lung may be more appropriate for measuring the host response.

Finally, our results compare the impact that culturing *S. aureus* in SCFM2 compared to agar plates has on both bacterial colonization and host response during *S. aureus* induced murine pneumonia. The agar plate preparation has been traditionally used in the lab to prepare bacteria for murine pneumonia infections (45–47). The choice of growing *S. aureus* in SCFM2 in wells was adapted from previous studies that used 4-well microchamber slides to study *S. aureus* transcriptomics, physiology, and spatial biogeography (28, 48). A 6-well plate format was chosen to maximize the surface area and increase the overall number of *S. aureus*. The comparison between the agar plate, our traditional bacterial method, and the SCFM2 preparation was adapted from a study from our group that compared the transcriptomics of *Pseudomonas aeruginosa* during murine pneumonia infections (49). When comparing growth kinetics of each strain in LB to SCFM2 in 6-well plates, no significant differences were observed between the two growth mediums. This indicates that the nutritional environment of the media did not significantly alter the growth kinetics of the bacteria and was not responsible for the differences observed in colonization. Based on our results, SCFM2 either had a neutral or beneficial impact on *S. aureus’* ability to colonize the lungs during acute pneumonia when compared to our standard laboratory protocol of culturing bacteria on agar plates. For WU1 in particular, we observed that growth in SCFM2 results in higher colonization in both the throat and lungs of Balb/c mice compared to agar plates. WU1’s ability to colonize the oropharynx has been described and our results indicate growth in SCFM2 improves colonization levels for 2 days post infection. WU1’s colonization in the lungs have not previous been explored and here we report that culturing WU1 in SCFM2 leads to a sustained long-term infection up to 4 days post infection. For all *S. aureus* strains tested here, while SCFM2 can improve lung colonization, the overall CFUs recovered from lungs were still lower than the number of originally administered bacteria. This suggest that while *S. aureus* is typically cleared by the host, growth in SCFM2 may improve the bacteria’s ability to persist beyond 24 hours in the lungs.

In conclusion, we have characterized the potential benefits culturing *S. aureus* in SCFM2 has on colonization during murine respiratory infections as well as the host response compared to LB media. While this study utilized six *S. aureus* strains, ultimately focusing on one, *S. aureus* is known to infect multiple human body site and how SCFM2 impacts isolates from different infection sites remains to be determined. Our study administered *S. aureus* to the mice intranasally but the effect of commonly used methods of administration, such as intratracheal, on the infection will be explored in future studies. Additionally, we report that WU1 infections in the lungs were well maintained up to 4 days post-infection and future studies will identify how long the infection can be maintained. The use of humanized mice has been shown to be more susceptible to *S. aureus* infections and allowed the study of human response to infection (50, 51). The infection outcome of humanized mice, such as CF humanized mice, when infected with *S. aureus* cultured in SCFM2 might further improve the clinical relevance of this infection model. Finally, while we observed that SCFM2 can increase colonization in the lungs, the mechanisms responsible are not understood. *In vivo* proteomics and transcriptomics might provide insight into the underlying benefits of SCFM2. Overall, our findings highlight the interplay between bacterial strain, mouse strain, and culture conditions on *S. aureus* ability to colonize during a murine pneumonia model and lays the groundwork for future studies.

## MATERIAL AND METHODS

### Bacterial Strains and Comparative Analysis

Six *S. aureus* strains were used in this study (Table 1). To determine the diversity of the *S. aureus* strains chosen, the whole genomes of each strain were compared bioinformatically. The sequence assemblies for JE2, Newman, N315, Sa_CFBR_43, and Sa_CFBR_46 was retrieved from NCBI. The whole genome sequence of WU1 was kindly provided by Dr. Dominique Missiakas (University of Chicago). All assemblies were compared to each other, and average nucleotide identity analysis was determined through pyani (52).

### Bacterial Growth Curves in LB and SCFM2 in 6-well Plates

Strains were streaked on *Staphylococcus* Isolation Agar (SIA) (lysogeny broth (LB) (BD) + 7.5% NaCl) and grown overnight at 37 . A single colony was transferred to a 2 ml of LB and grown shaking at 37 . The cultures were then transferred to a 1.5 ml Eppendorf tube, washed 3 times with sterile 1X phosphate buffer saline (PBS), and optical density was measured at 600 nm (OD_600_). The cultures were adjusted to an OD_600_ of 0.01 in 6 ml of LB or SCFM2 (Synthbiome) and were and grown statically in a 6-well plate (Cellstar) at 37 . Aliquots of each culture were removed every 2 hours for 8 hours as well as 24 hours, serially diluted, and plated on Lysogeny agar (LA) plates. The plates were placed in at 37 and the number of colony-forming units (CFUs) was determined the following day.

### *S. aureus* Growth on Agar Media or SCFM2 Workflow

To prepare the *S. aureus* inoculum under our agar plate procedure, strains were streaked on SIA and grown overnight at 37 . On the following day a small swab of each *S. aureus* was transferred into a sterile Eppendorf tube containing sterile 1X PBS. The strains were washed 3 times and OD_600_ was measured. Cultures were adjusted to an OD_600_ of 6.5 in 1 ml of 1X PBS, corresponding to ∼1 x 10^8^ colony-forming units (CFU) per 25 µl dose. An aliquot was removed, serially diluted, and plated on SIA plates to determine colony forming units per dose.

For *S. aureus* inoculum preparation in SCFM2, strains were streaked on SIA and grown overnight at 37 . A single colony was transferred to a culture tube with 2 ml of LB and grown shaking at 37 overnight. The cultures were then pelleted, washed 3 times with 1X PBS, and the OD_600_ was measured. The strains were then adjusted to an OD_600_ of 0.01 in 6 ml of SCFM2 in a 6-well plate and were grown statically at 37 for 24 hours. After incubation, the cultures were transferred to a 15 ml falcon tube, pelleted, and concentrated 15X in fresh SCFM2. An aliquot was removed, serially diluted, and plated on SIA plates to determine CFU per dose.

### Mice

All animal experiments were performed according to the guidelines of the Emory University Institutional Animal Care and Use Committee under the approved protocol PROTO201700441. 8-to 10-week-old female C57BL/6 and Balb/c mice were purchased from Jackson Laboratories (Bar Harbor, ME) and were acclimated for at least a week at Emory University before experiments. Prior to infection, all mice were anesthetized with an intraperitoneal injection of a 0.2 ml mixture of ketamine (6.7 mg/ml) and xylazine (1.3 mg/ml). All mice were euthanized by CO_2_ asphyxiation.

### Lung Infection

Anesthetized C57BL/6 and Balb/c mice were intranasally administered 25 µl (12.5 µl per nostril) of each strain cultured either on agar plates or in SCFM2 preparation corresponding to about 1 x 10^8^ CFUs. Mice were humanely euthanized at specified time points and whole lungs and nasal wash were aseptically collected. For the nasal wash, 1 ml of sterile 1X PBS was flushed through the nasal passage using an 18-G catheter placed at the nasopharyngeal opening. The lungs were weighed and homogenized in 1 ml of sterile 1X PBS in a bullet blender storm 5 (Next Advance). Both the nasal wash and lungs were serially diluted and plated on SIA. Plates were then incubated at 37 overnight and CFUs were determined.

### Oropharyngeal Colonization

Anesthetized Balb/c mice were intranasally administered 25 µl (12.5 µl per nostril) of WU1 prepared either through the agar plate or SCFM2 workflow. To track WU1 colonization in the throat over time, mice were anesthetized with 3% isoflurane through an XGI-8 Gas Anesthesia System (Caliper Life Sciences) and oropharyngeal swabs were collected from each mouse using calcium alginate swabs (Puritan) on days 1, 2, 3, 7, 10, and 14. The swab was cut from the stick with sterile scissors, transferred to a 1.5 ml Eppendorf tube with 500 µl PBS, resuspended, serially diluted, and spread on SIA plates. The plates were then incubated at 37 overnight and CFUs were determined.

### Lung Preparation for Histology and Proinflammatory Cytokine Analysis

To prepare the left or right lung lobes for histology and cytokine analysis, anesthetized Balb/c mice were intranasally administered 25 µl (12.5 µl per nostril) of *S. aureus* strain WU1 prepared either through the agar plate or SCFM2 workflow. 1X PBS and SCFM2 were also administered as a “vehicle control”. After 24 hours, mice were euthanized, and the chest was opened. A sterile clamp was used to separate the lung lobes, one lobe was inflated with 4% paraformaldehyde, and removed for histology. The other lobe was processed for cytokine analysis. To determine the bacterial loads in each lung, a parallel group of mice (n=2) were used. Each lobe was separated, weighed, and homogenized in 1 ml of sterile 1X PBS. The lung lobes were plated on SIA, incubated overnight at 37, and CFUs were counted.

### Histopathology

Lung specimen from mice were insufflated with, and fixed in, 4% paraformaldehyde (PFA), processed, and blocked in paraffin for histological analysis. All samples were sectioned at 5 µm and stained with hematoxylin-eosin (H&E) for routine histopathology. Samples were evaluated by a board-certified veterinary pathologist in a blinded manner. Sections were examined under light microscopy using an Olympus BX51 microscope and photographs were taken using an Olympus DP73 camera.

### Cytokine Analysis

Lung lobes were weighed, homogenized in 1 ml of sterile 1X PBS supplemented with ProBlock Protease Inhibitor Cocktail (Goldbio), the supernatant was collected through centrifugation, and stored at -80 . Cytokine levels were measured through the MesoScale Discovery (MSD) PlatformV-PLEX proinflammatory Panel 1 Mouse Kit. The cytokines measured through this kit were IFN-⍰, IL-1β, IL-2, IL-4, IL-5, IL-6, IL-10, IL-12p70, KC/GRO, and TNF-11. The cytokine levels were quantified and analyzed at the Emory Multiplex Immunoassay Core with MESO QuickPlex SQ 120 following the manufacturer’s instructions. Results were normalized to lung weight. Results for IL-4 were not included as the results from all groups were below the limit of detection.

### Statistical Analysis

Statistical analyses were performed for all experiments using GraphPad Prism 9.

## ACKNOWLEDGEMENTS

We thank Dr. Dominique Missiakas for providing the *S. aureus* WU1 strain as well as its associated genomic sequences. Additionally, the N315 strain was kindly provided by Dr. Timothy Read. We thank Dr. Christine Bojanowski for the fruitful discussions. We also thank the Emory Histology and Molecular Pathology Lab as well as the Emory Multiplexed Immunoassay Core for their help with performing the histopathology and cytokine analysis, respectively. Finally, we thank the Goldberg Lab for their helpful discussion and feedback on the manuscript.

This work was supported by Cystic Fibrosis Foundation Grants WHITEL20A0 and WHITEL24XX0. The pathology research reported in this publication was supported by the Emory National Primate Research Center of the Office of Research Infrastructure Programs/OD P51 OD011132. The content is solely the responsibility of the authors and does not necessarily represent the official views of the National Institutes of Health.

