## Supplemental Material for "Growing *Staphylococcus aureus* in Synthetic Cystic Fibrosis Medium Promotes Colonization in a Murine Pneumonia Model"

**Table S1. Average nucleotide identity values.**

| Strain | JE2 | N315 | Newman | Sa_CFBR_43 | Sa_CFBR_46 | WU1 |
| --- | --- | --- | --- | --- | --- | --- |
| JE2 | 1 | 0.98950869 | 0.999075822 | 0.99987798 | 0.973625079 | 0.990052994 |
| N315 | 0.989491561 | 1 | 0.9894661 | 0.989440433 | 0.973406043 | 0.989826162 |
| Newman | 0.999024674 | 0.989454179 | 1 | 0.99848061 | 0.973404448 | 0.99000707 |
| Sa_CFBR_43 | 0.99989204 | 0.98936415 | 0.998515885 | 1 | 0.973239822 | 0.990193949 |
| Sa_CFBR_46 | 0.973552667 | 0.973405941 | 0.973365271 | 0.973253801 | 1 | 0.973250535 |
| WU1 | 0.99007895 | 0.989852571 | 0.990038341 | 0.990103116 | 0.973251346 | 1 |

Average nucleotide identity was determined between the genomic sequences of each strain using pyani. The table depicts the numerical value outputs from the analysis.

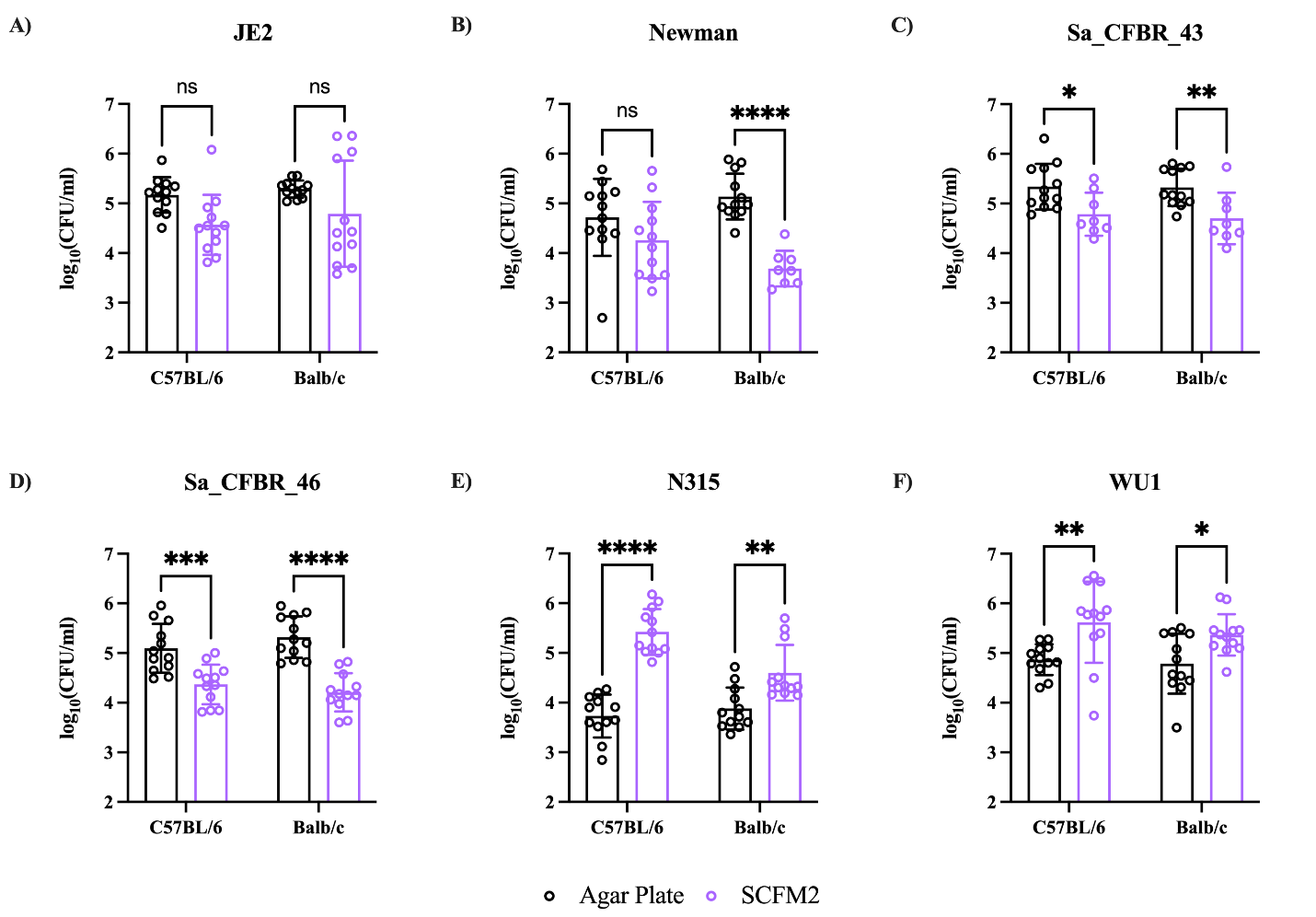

**Supplemental Figure 1. Impact of different culture conditions on nasal cavity colonization in mice during acute pneumonia.**

*S. aureus* strains were cultured either on agar plates or in SCFM2 as described in Figure 3 and administered to either 8–10-week-old female C57BL/6 and Balb/c mice. Mice were sacrificed after 24 hours post-infection, 1 ml of sterile 1X PBS was flushed through the nasal passage using an 18-G catheter placed at the nasopharyngeal opening. The washes were serially diluted and plated on SIA. Each data point represents a mouse. Data was analyzed using two-way ANOVA with Šídák correction. The mean and standard deviation are represented by the error bars. *p < 0.05, **p < 0.01, ***p < 0.001, ****p < 0.0001, ns=not significant.

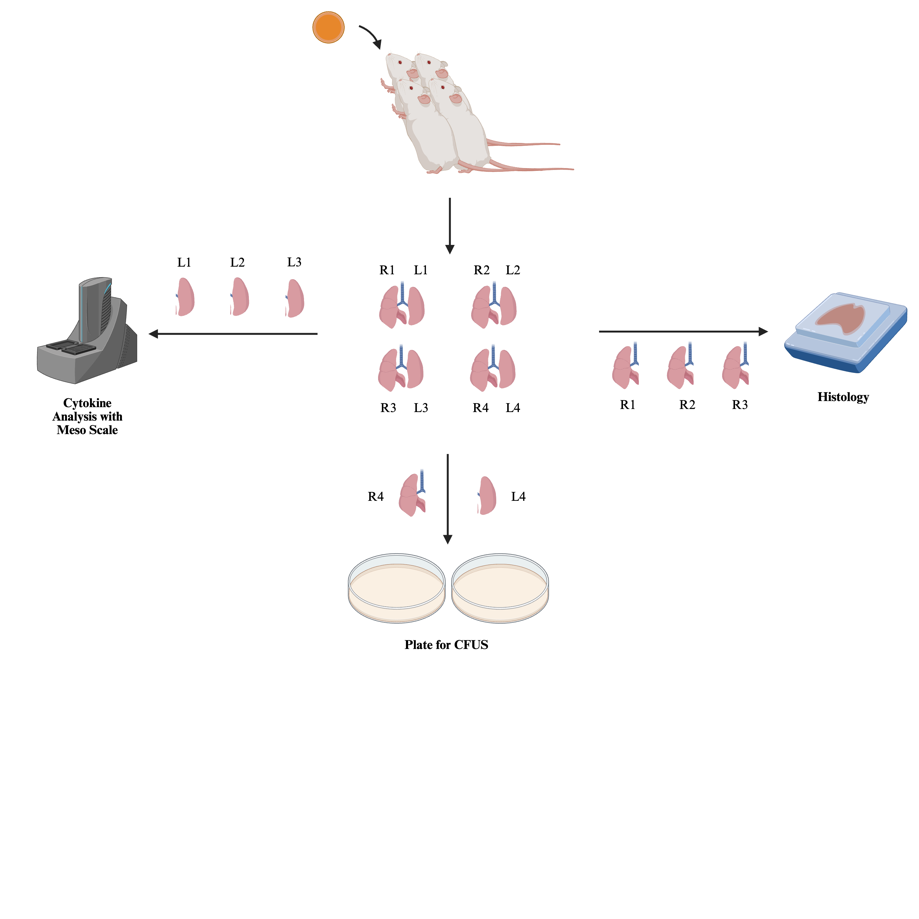
**A)**

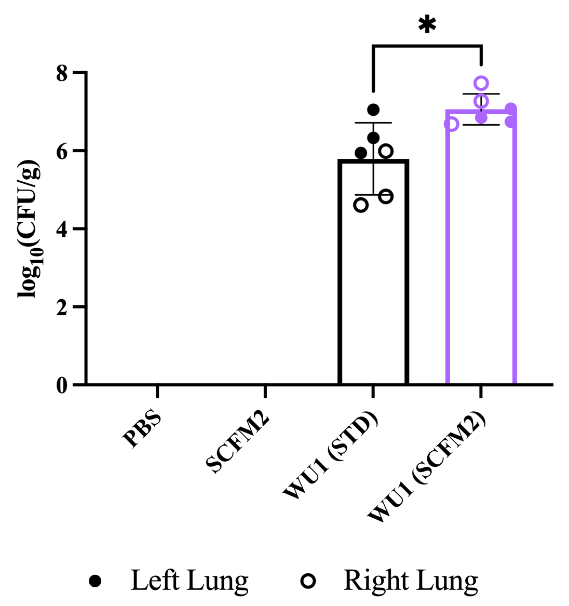

**B)**

**Supplemental Figure 2. Workflow for lung lobe separation for cytokine analysis and histology.**

**A)** Seven 8–10-week-old Balb/c mice were used for each condition. Mice were sacrificed 24 hours post infection and for each mouse, a clamp was inserted to separate the left and right lobes. For 3 mice, one lobe was inflated with 4% paraformaldehyde, separated, and stored in 4% paraformaldehyde for histology. The other lung lobe was homogenized in 1X PBS + 1X protease inhibitor, the homogenate supernatant was isolated, and cytokines were measured with the V-plex proinflammatory cytokine panel 1 (MSD). In 3 other mice, the alternate lung lobe was process as described above leading to 3 left and 3 right lung lobes from different mice for histology and cytokine analysis. **B)** The left and right lobe from the 7^th^ mouse was used to determine CFUs as described above. Statistical analysis was performed using two-way ANOVA with Fisher’s LSD post-hoc test. *p < 0.05.
